# Closing the loop: Teaching single-cell foundation models to learn from perturbations

**DOI:** 10.1101/2025.07.08.663754

**Authors:** Yash Pershad, Tarak N Nandi, Joseph C Van Amburg, Alyssa C Parker, Luiza Ostrowski, Hannah K Giannini, David Ong, J Brett Heimlich, Esther A Obeng, Katrin Ericson, Anupriya Agarwal, Ravi K Madduri, Alexander G Bick

**Affiliations:** Department of Medicine, Vanderbilt University Medical Center, Nashville, TN, USA; Vanderbilt Genetics Institute, Vanderbilt University, Nashville, TN, USA; Data Science and Learning, Argonne National Laboratory, Lemont, IL, USA; Consortium for Advanced Science and Engineering, University of Chicago, Chicago, IL, USA; Knight Cancer Institute, Oregon Health & Science University, Portland, OR, USA; Department of Oncology, St. Jude Children’s Research Hospital, Memphis, TN, USA; RUNX1 Research Program, Santa Barbara, California, USA

## Abstract

The application of transfer learning models to large scale single-cell datasets has enabled the development of single-cell foundation models (scFMs) that can predict cellular responses to perturbations in silico. Although these predictions can be experimentally tested, current scFMs are unable to “close the loop” and learn from these experiments to create better predictions. Here, we introduce a “closed-loop” framework that extends the scFM by incorporating perturbation data during model fine-tuning. Our closed-loop model improves prediction accuracy, increasing positive predictive value in the setting of T-cell activation three-fold. We applied this model to RUNX1-familial platelet disorder, a rare pediatric blood disorder and identified two therapeutic targets (mTOR and CD74-MIF signaling axis) and two novel pathways (protein kinase C and phosphoinositide 3-kinase). This work establishes that iterative incorporation of experimental data to foundation models enhances biological predictions, representing a crucial step toward realizing the promise of "virtual cell" models for biomedical discovery.

## Introduction

Predicting how a cell will respond to a perturbation is a significant unsolved biological challenge that would advance our understanding of biological mechanisms and therapeutic development.^1–3^ Recent progress in artificial intelligence has led to the development of single-cell foundation models (scFMs) – deep learning models which are pre-trained on vast amounts of single-cell data and can be fine-tuned for specific tasks. scFMs represent an important step toward creating "virtual cells" that can simulate cellular responses to diverse perturbations without requiring extensive experimental validation.^2,4–8^ Specifically, scFMs enable *in silico* perturbation (ISP), where a fine-tuned model predicts how cell states would shift in response to a genetic perturbations (e.g., knockout or over-expression of a gene).^1,2,5,6^ This approach holds particular value for rare diseases, where patient samples are scarce so experimental screening with patient samples is challenging. Moreover, the consequences of rare genetic mutations often manifest in specific cell lineages that may be inaccessible to sampling or experimental perturbation.

Models of “virtual cells” can help overcome these limitations by simulating cellular states *in silico* and prioritizing interventions most likely to restore normal cellular function.^2^ These models generate a set of predictions which can be experimentally validated. Ideally, the results of these experiments would be used to improve the model, thereby “closing the loop” between computational prediction and experimental evaluation.^9^ Despite their promise, the accuracy of “open-loop” ISP predictions remains poorly characterized and “closed-loop” models which leverage observed perturbation data to improve ISP predictions do not exist.

Here, we develop and systematically evaluate ISP performance using a state-of-the-art scFM Geneformer-30M-12L^5^ and introduce a “closed-loop” ISP framework that extends the scFM by incorporating experimental perturbation data during model fine-tuning. We demonstrate the utility of this approach in two distinct human biological settings: (1) T-cell activation, a widely studied problem with applications in cancer immunotherapy, autoimmunity, and infectious disease, and (2) *RUNX1*-familial platelet disorder (*RUNX1*-FPD), a rare hematologic disorder affecting approximately 20,000 people in the US that predisposes patients to early-onset myeloid leukemia.^10–12^ Our closed-loop approach significantly improves prediction accuracy, increasing positive predictive value three-fold while maintaining high negative predictive value and enhancing both sensitivity and specificity. This framework identifies and validates multiple therapeutic targets for *RUNX1*-FPD, demonstrating its potential for accelerating rare disease drug discovery.

## Results

### Benchmarking open-loop in silico perturbation predictions

We evaluated open-loop *in silico* perturbation (ISP) predictions using the Geneformer-30M-12L single-cell foundation model.^5,13^ For systematic evaluation, we utilized existing CRISPRi and CRISPRa screens of over 18,000 genes that measured IL-2 and IFN-γ production as a marker of activation after CD3-CD28 stimulation^14^, providing an orthogonal modality from transcriptomics for assessing genetic perturbation effects on T cell activation.

We fine-tuned Geneformer-30M-12L to predict T cell activation status using data from four studies where T cells were stimulated via CD3-CD28 beads or phorbol myristate acetate/ionomycin (PMA/ionomycin).^15–18^ While the pre-trained model embeddings clustered by study rather than activation status, the fine-tuned model successfully classified cells by activation status, with an accuracy of 99.8% and macroF1 of 0.998 on a hold-out test set of cells (**Figure S1**). Using this fine-tuned model, we performed ISP across 13,161 genes, simulating both gene overexpression and knockout to model CRISPRa and CRISPRi respectively, and validated predictions against flow cytometry data (**Figures 1A-B**). We found that for T-cell activation open-loop ISP predictions from Geneformer-30M-12L and Geneformer-106M-12L were not significantly different (Spearman’s ρ = 0.90), so we subsequently used the lighter weight model.

**Figure 1.**
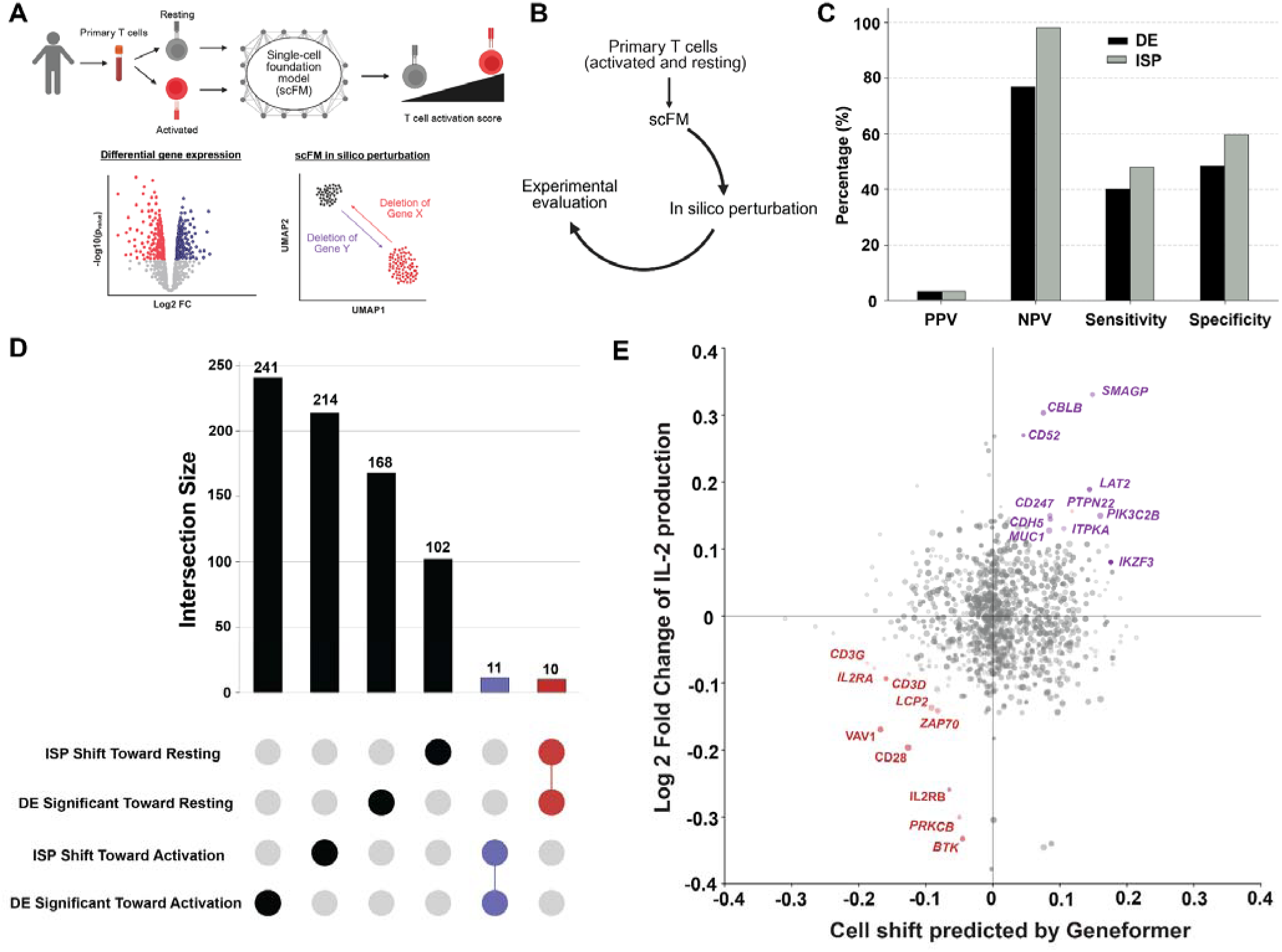
We apply open-loop *in silico* perturbation to primary human T-cell activation. **A)** Fine-tuning a classifier to predict T cell activation status using data from four studies where primary T cells were stimulated via CD3-CD28 beads or phorbol myristate acetate/ionomycin (PMA/ionomycin). We compared predictions from differential expression (DE) and single-cell foundation model (scFM) in silico perturbation (ISP). **B)** An “open-loop” approach, where the model fine-tuned using primary T cells which are resting or activated makes ISP predictions about what genetic perturbations would alter a T cell’s state and shift the cell toward resting or stimulated. For systematic evaluation, we utilized Schmidt et al’s CRISPRi and CRISPRa screens of over 18,000 genes that measured IL-2 and IFN-γ production as a marker of activation after CD3-CD28 stimulation, providing an orthogonal modality from transcriptomics for assessing genetic perturbation effects on T cell activation.^14^ **C)** Performance metrics comparing DE and ISP approaches against the flow cytometry experimental data from Schmidt et al.^14^ Both methods achieve ∼3% positive predictive value (PPV), while ISP shows superior negative predictive value (NPV; 98% vs 78%), sensitivity (48% vs 40%), and specificity (60% vs 50%). **D)** Intersection analysis of significant predictions from ISP and DE methods. Numbers indicate predicted genes shifting T cells toward resting (top) or activation (bottom) states. Most predictions are method-specific, with only 21 genes identified by both approaches. Purple set represents the shared set between DE and ISP predicted to shift the cells toward activation. Red set represents the shared set between DE and ISP predicted to shift the cells toward resting. **E)** Scatter plot showing predicted cell state shifts by ISP (x-axis) versus actual IL-2 production changes from flow cytometry (y-axis). Key T cell activation genes (red) and inactivation genes (purple) are correctly predicted to shift cells toward resting state upon knockout.

When comparing performance metrics using flow cytometry as ground truth, both open-loop ISP and differential expression (DE) achieved positive predictive values (PPVs) of 3%. However, open-loop ISP demonstrated superior performance compared to differential expression (DE) for negative predictive value (98% versus 78%), sensitivity (48% versus 40%), and specificity (60% versus 50%) (**Figure 1C**). Notably, genes identified as significant by both DE and open-loop ISP showed an improved PPV of 7% (**Figure 1C**). These results indicate that open-loop ISP performs at least equivalently to DE, the current gold standard for identifying disease-relevant gene expression changes, while offering improved ability to correctly identify true negatives.

Despite using the same underlying data, only 2.9% of predictions from open-loop ISP and DE were overlapping (**Figure 1D**). Both methods predicted more genes shifting T cells toward activation than toward a resting state. Only 21 genes were predicted by both methods to have effects in the same direction: 11 shifting toward activation and 10 toward resting (**Supplemental Table 1**). While predictions from either method alone exhibit low PPV, gene sets predicted by both ISP and DE show higher positive predictive value and represent key T cell activation genes which, when knocked out, would shift cells toward a resting state, including *IL2RA*, *VAV1*, *ZAP70*, *CD3D*, *CD3G*, and *LCP2* (**Figure 1E**).

### Closing the loop: Incorporation of perturbation examples in fine-tuning improves prediction accuracy

We next sought to improve the PPV of open-loop ISP. We hypothesized that incorporating experimental perturbation data into the fine-tuning of Geneformer-30M-12L could enhance ISP performance – a process we termed "closed-loop" ISP (**Figure 2A**), wherein the closed-loop refers to the iterative process of model refinement using feedback from experimental data to guide fine-tuning. We fine-tuned the model with single-cell RNA sequencing (scRNAseq) data from CRISPR activation and interference screens in primary human T cells (i.e., Perturb-seq) alongside our existing scRNAseq data from resting and activated T cells. Critically, the Perturb-seq data was labeled only with activation status, not with which gene was perturbed. We then performed ISP using this new fine-tuned model on all genes except the 75 perturbed genes in the screens (**Supplemental Table 2**).

**Figure 2.**
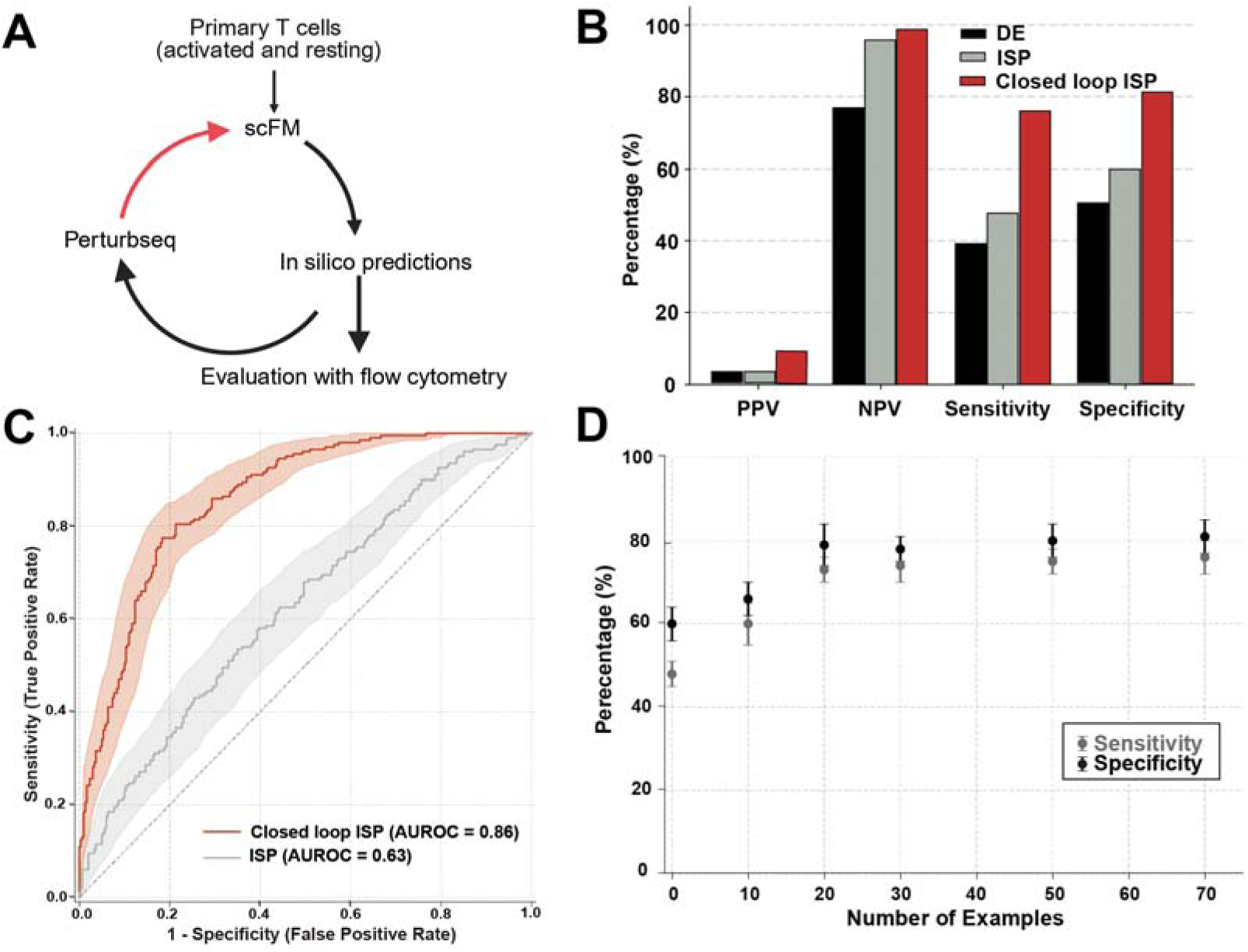
Closed-loop *in silico* perturbation improves prediction accuracy through incorporation of experimental perturbation data. **A)** Schematic of closed-loop ISP framework. Primary T cells in resting and activated states are used to fine-tune the scFM, with additional incorporation of Perturb-seq data creating a feedback loop between computational predictions and experimental validation. **B)** Performance comparison of standard DE, ISP, and closed-loop ISP. Closed-loop ISP achieves three-fold improvement in PPV (9% vs 3%) while maintaining high NPV (99%) and improving sensitivity (76%) and specificity (81%). **C)** Receiver operating characteristic (ROC) curves comparing ISP and closed-loop ISP performance. Closed-loop ISP achieves significantly higher AUROC (0.86; 95% CI: 0.83-0.89) compared to standard ISP (0.63; 95% CI: 0.58-0.68). **D)** Learning curve showing model performance as a function of perturbation examples used in fine-tuning. Sensitivity and specificity improve dramatically with 10 examples and plateau at approximately 20 examples. Error bars represent standard error across different random subsets of perturbation examples.

Closed-loop ISP increased PPV three-fold—from 3% to 9%—with concurrent improvements in negative predictive value (99%), sensitivity (76%), and specificity (81%) (**Figure 2B; Supplemental Table 3**). The area under the receiver operator characteristic curve (AUROC) was significantly higher for closed-loop ISP (0.86; 95% confidence interval: 0.83-0.89) compared to standard ISP (0.63; 95% CI: 0.58-0.68).

To determine the minimum number of perturbation examples required for substantial improvement, we evaluated model performance with incremental additions of random perturbation subsets. Performance metrics improved dramatically with just 10 examples (sensitivity: 61%, 95% CI: 58% to 64%; specificity: 66%, 95% CI: 62% to 70%) and approached saturation at approximately 20 examples (sensitivity: 76%, 95% CI: 72 to 78 specificity: 79%, 95% CI: 75% to 83%) (**Supplemental Table 4**). After this point, performance did not improve significantly with additional examples (**Figure 2D**). Standard errors of sensitivity and specificity ranged from 5% to 10%, indicating that the specific perturbation examples used in fine-tuning significantly influence model predictions. These findings suggest that even a modest number of experimental validations can substantially enhance closed-loop ISP accuracy compared to baseline ISP.

### In silico perturbation prioritizes gene targets for RUNX1-Familial Platelet Disorder

To demonstrate the utility of this closed-loop virtual cell model, we next applied the closed-loop framework to *RUNX1*-familial platelet disorder (*RUNX1*-FPD), a rare pediatric onset hematologic disease affecting >18,000 people in the United States.^10–12^ *RUNX1*-FPD is caused by loss-of-function mutations in *RUNX1*, affecting hematopoietic stem cells (HSCs) and characterized by thrombocytopenia, impaired platelet function, immune dysregulation, and increased risk of early-onset myeloid neoplasms.^19–21^ Currently, no interventions exist to prevent progression to myeloid neoplasms.

Since patient samples are scarce, we leveraged human HSCs engineered to have *RUNX1* loss of function mutations that model *RUNX1*-FPD.^22,23^ We generated scRNAseq data from these *RUNX1* engineered HSCs (**Figure S2**). We first validated that the engineered cells with scRNAseq data from 10 *RUNX1*-FPD patients.^24^ We found high concordance between the *RUNX1*-engineered HSCs and *RUNX1*-FPD patient HSCs; for example, the downstream targets of RUNX1 had reduced expression in patient HSCs also had reduced expression in our human *RUNX1*-FPD HSC model (**Figure S3**).

We fine-tuned Geneformer-30M-12L to classify HSCs between *RUNX1*-engineered HSCs and control HSCs whose guide RNA targeted *AAVS1* (**Figure 3A**). The fine-tuned model successfully distinguished these two cell states (**Figure S4**). Using this fine-tuned model, we performed open-loop ISP to identify genes that, when deleted, would shift *RUNX1*-knockout HSCs toward a control-like state (**Figure 3B; Supplemental Table 5**). Comparing DE and ISP results, we identified 14 genes predicted by both methods to significantly shift *RUNX1*-knockout cells toward control cells (**Figure 3C**). From these targets, we selected eight genes with available specific small molecule inhibitors: *PRKCB*, *UBB*, *CHKA*, *CAMK2G*, *HSPA8*, *MDM4*, *YBX1*, and *PIK3C2B*.

**Figure 3.**
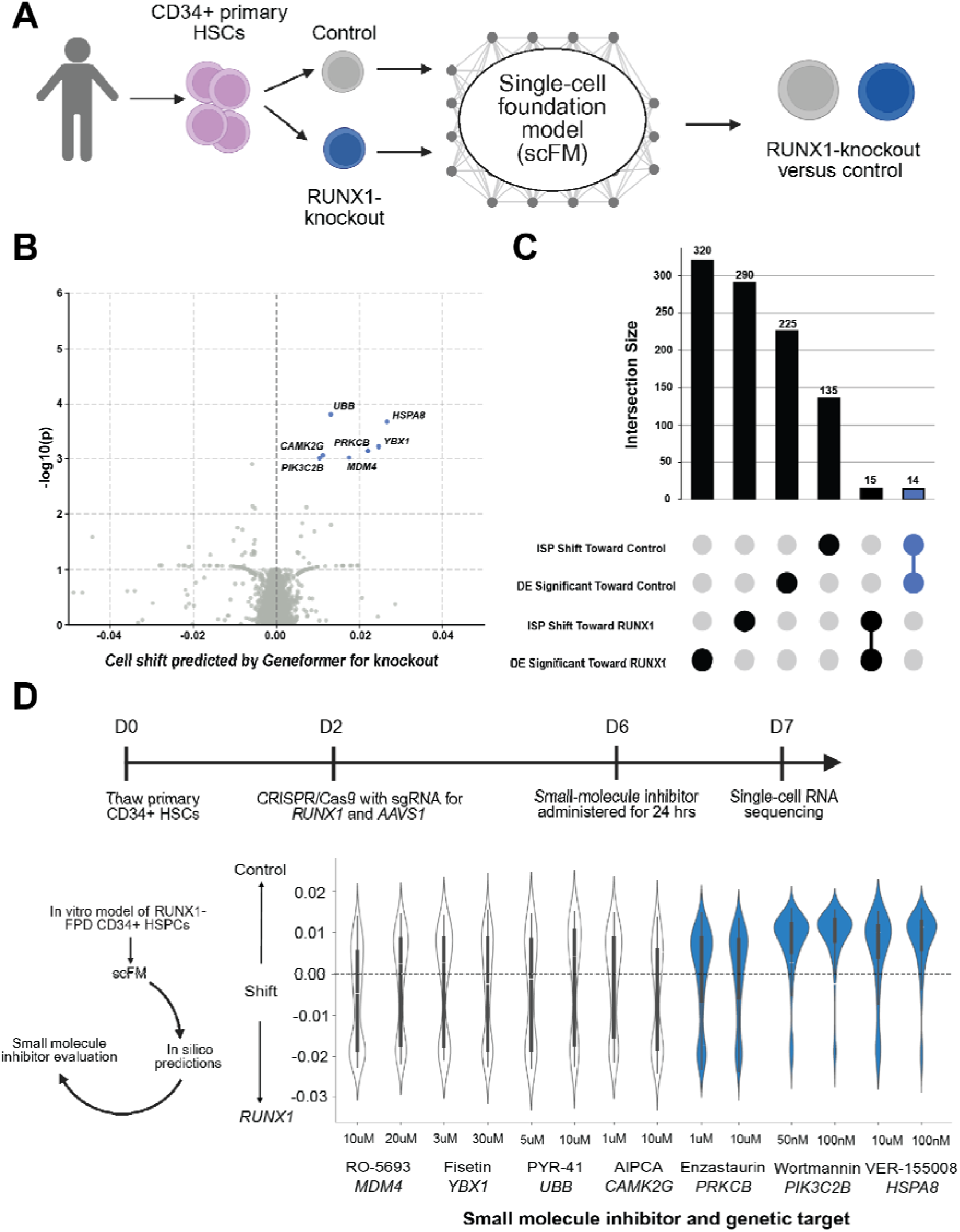
Open-loop *in silico* perturbation identifies a set of therapeutic targets for *RUNX1*-familial platelet disorder. **A)** Experimental design for *RUNX1*-FPD target discovery. Primary CD34+ hematopoietic stem cells (HSCs) are edited with CRISPR/Cas9 targeting *RUNX1* or *AAVS1* (control), followed by single-cell RNA sequencing and fine-tuning of the scFM to classify cells by guide RNA. **B)** Volcano plot of ISP predictions showing genes that shift *RUNX1*-knockout cells toward control state when deleted. Eight genes with available small molecule inhibitors are highlighted (*UBB*, *HSPA8*, *CAMK2G*, *PRKCB*, *YBX1*, *PIK3C2B*, *MDM4*, *CHKA*). **C)** Intersection analysis of predictions from ISP and DE methods. Of 15 genes predicted by both methods, 14 shift cells toward the control state (blue), while one shifts toward the *RUNX1*-knockout state (grey). **D)** Experimental validation workflow and results. CD34+ HSCs are edited with *RUNX1* or *AAVS1* sgRNAs, treated with small molecule inhibitors, and analyzed by single-cell RNA sequencing. Violin plots show the distribution of cell embedding shifts toward control centroids (positive shift) or toward *RUNX1*-knockout centroids (negative shift) for each treatment. Three inhibitors demonstrate significant phenotypic rescue (blue): enzastaurin (PRKCB inhibitor), wortmannin (PIK3C2B inhibitor), and VER-155008 (HSPA8 inhibitor).

To experimentally validate our predictions, we evaluated the effect of these inhibitors on cell state using scRNAseq. After CRISPR targeting *RUNX1* and *AAVS1*, we treated progenitor cells with small molecule inhibitors for 24 hours and performed scRNAseq (**Figure 3D**). By analyzing cell embeddings, we tested whether chemically perturbed cells shifted significantly from the *RUNX1* centroid toward the control centroid. Three of eight inhibitors demonstrated significant rescue of the *RUNX1*-knockout phenotype: enzastaurin (*PRKCB* inhibitor), wortmannin (*PIK3C2B* inhibitor), and VER-155008 (*HSPA8* inhibitor). These phenotypic shifts increased with higher doses and were specific to *RUNX1*-FPD cells (**Figure 3D**).

### Closed-loop in silico perturbation identifies therapeutic targets for RUNX1-FPD

We next applied the closed-loop ISP framework to *RUNX1*-FPD. Since our approach does not require annotation of which genes were perturbed, we could utilize chemical perturbations, which are more convenient and cost-effective than genetic perturbations. We performed a chemical perturbation screen using 35 small-molecule inhibitors administered to *RUNX1*-knockout HSCs, followed by scRNAseq (**Figure 4A**; **Supplemental Table 6**). We then performed ISP with this fine-tuned model on all genes except the genetic targets of the 35 drugs included in the screen (**Figure S5**). After multiple hypothesis correction, closed-loop ISP predicted 6 targets for inhibition: *CD74*, *JAK1*, *JAK2*, *FKBP1A*, *PIK3C2B*, and *PRKCB* (**Figure 4B; Supplemental Table 5**).

**Figure 4.**
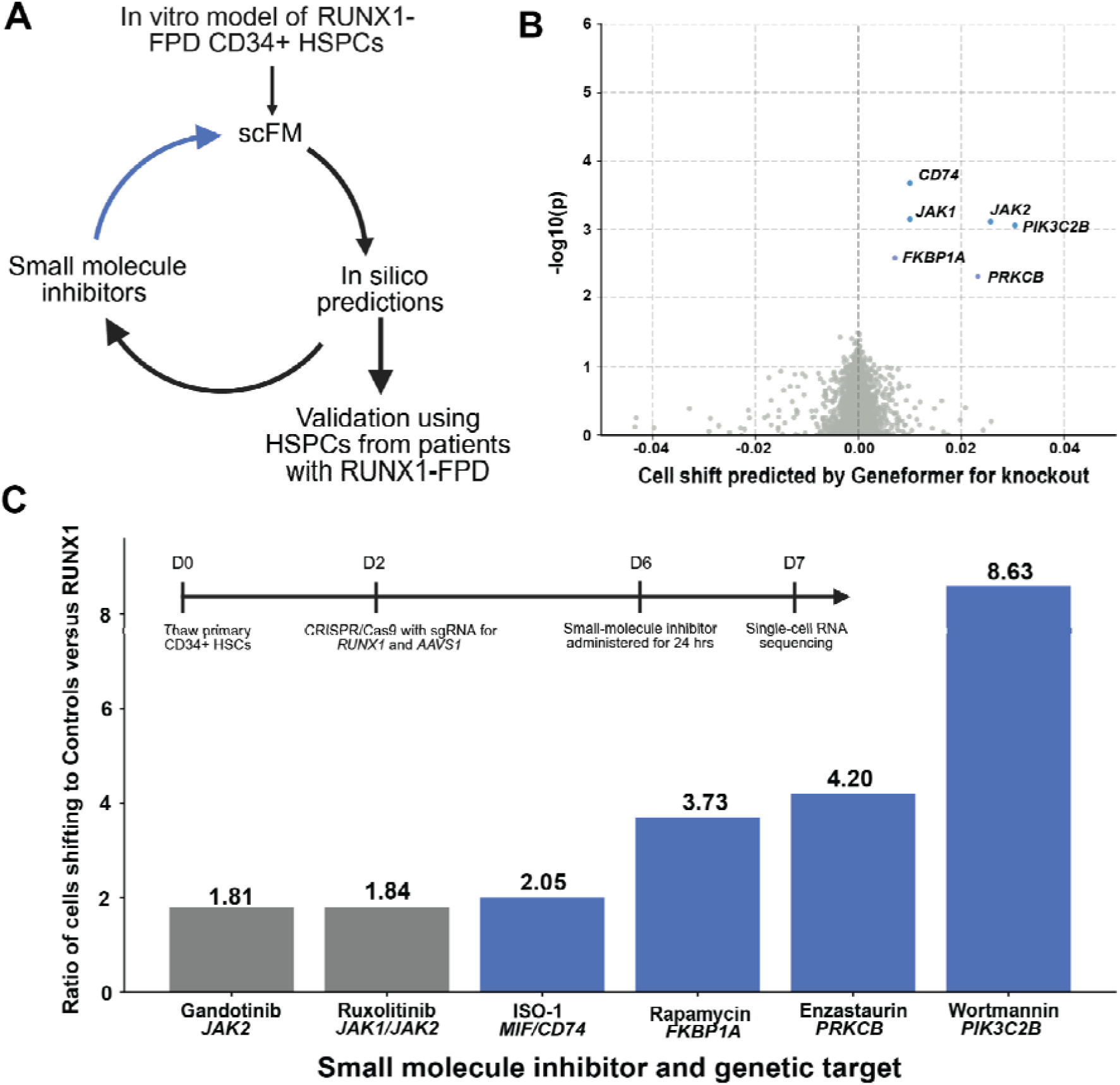
Closed-loop in silico perturbation identifies validated and novel therapeutic targets for *RUNX1*-FPD. **a)** Closed-loop ISP framework applied to *RUNX1*-FPD. Small molecule inhibitors are administered to the *in vitro RUNX1*-FPD model, followed by single-cell RNA sequencing and ISP predictions, which are then validated in HSCs from *RUNX1*-FPD patients. **b)** Volcano plot showing closed-loop ISP predictions. Six targets are significant after multiple hypothesis correction: *CD74*, *JAK1*, *JAK2*, *FKBP1A*, *PIK3C2B*, and *PRKCB*. **c)** Validation of predicted targets. Primary CD34+ HSCs are edited with *RUNX1* or *AAVS1* sgRNAs, treated with inhibitors targeting predicted genes, and analyzed by single-cell RNA sequencing. Bar chart shows ratio of cells shifting toward control versus *RUNX1*-knockout state. Wortmannin (*PIK3C2B* inhibitor) shows strongest effect (8.63), followed by enzastaurin (*PRKCB* inhibitor, 4.20), rapamycin (*FKBP1A*/*mTOR* inhibitor, 3.73), and ISO-1 (*CD74*-*MIF* interaction inhibitor, 2.05). Grey bars indicate inhibitors with minimal effect; blue bars indicate significant phenotypic rescue.

To validate these predictions, we administered small molecule inhibitors targeting these genes to HSCs in our *in vitro RUNX1*-FPD model and performed single-cell RNA sequencing. We evaluated whether treated cells shifted significantly from the *RUNX1*-knockout centroid toward the *AAVS1* control centroid and calculated the ratio of cells that shifted toward the control state versus those that did not shift or shifted toward *RUNX1*-knockout. Wortmannin (PI-3 kinase inhibitor targeting *PIK3C2B*) demonstrated the most significant shift, followed by enzastaurin (protein kinase C inhibitor targeting *PRKCB*), rapamycin (mTOR inhibitor targeting *FKBP1A*), and ISO-1 (inhibitor of CD74-MIF interactions). The genes identified in ISP as shifting *RUNX1*-knockout cells to a control state were significantly enriched for the MSigDB pathways involving PI3K/AKT/mTOR (adjusted p-value = 0.01).

## Discussion

In sum, for the first time, we systematically benchmark predictions from a scFM and show that “closing the loop” – that is, fine-tuning scFMs with strategic genetic and chemical perturbation data – sufficiently enhances accuracy to make experimental validation of its predictions tractable. Our work has several important implications for the development of virtual cell models.

First, we demonstrate the concrete value of “closing the loop” in building better virtual cell models. The concept of deep learning models iteratively improving using observed example data have been employed to build better models in a variety of applications, including antibody development, chemical discovery, autonomous vehicles, and robotics.^25–29^ Some have speculated that this strategy would also apply to single-cell biology.^2^ Here, we demonstrate that predictions from scFMs at baseline are susceptible to a high false positive rate, which can be significantly improved through “closing the loop”. In the case of T-cell activation, only 3% of predictions were true positives, which corroborates prior work that *in silico* perturbation without “closing the loop” is similar to performance by linear models.^30,31^ By closing the loop, with even a modest amount of perturbation data (e.g., 20 examples for T cell activation) increases PPV by three-fold (from 3% to 9%). These results highlight the utility of ongoing efforts to generate large-scale perturbation data in primary cells as well as the promise of future more advanced “closed-loop” models to incorporate such data.^14,32–34^

Second, we demonstrate the potential of virtual cell models for rare disease drug discovery. Two of the targets identified by our closed-loop ISP virtual cell model have strong biological evidence and are under investigation for *RUNX1*-FPD treatment. Notably, our team has recently launched a phase 2 clinical trial of low-dose sirolimus, an mTOR inhibitor, that was identified by our closed-loop ISP, to investigate safety and tolerability, with secondary endpoints investigating platelet count and function along with dynamics of premalignant clonal hematopoiesis clones (NCT06261060). Moreover, recent *in vivo* work has identified the CD74-MIF signaling axis as a targetable node in *RUNX1*-FPD to reduce inflammation and restore erythroid differentiation.^24^ Further, the closed-loop ISP framework established here can be readily extended to other diseases where single-cell data are available and is particularly valuable for rare diseases where patient data is scarce and comprehensive experimental perturbation studies are impractical.

Our study has several limitations. First, we benchmarked the closed-loop performance improvement using only a single scFM, Geneformer-30M-12L. We selected this model because it readily enables incorporation of perturbation examples in its ISP framework via a fine-tuned classifier, it has a similar architecture and performance to a range of other contemporary scFMs, and its ISP predictions were very similar to that of the larger Geneformer-106M-12L.^4,6,7,13^ Second, we comprehensively benchmarked the performance in the context of T cell activation, due to the availability of validation data. How this performance generalizes across a range of biological contexts will require further exploration. Third, our “virtual cell model” only enables examining the effects of *in silico* perturbation in one cell type at a time, so our experimental work which focused on HSCs represents only one aspect of *RUNX1*-FPD cell biology.

Overall, our work establishes that “closing the loop” creates better virtual cell models. This advance is a prerequisite to creating autonomous biological laboratories which can make, evaluate, and iteratively improve predictions without human supervision. Such laboratories have the potential to accelerate the pace of biomedical research.

---

**Figure S1.**
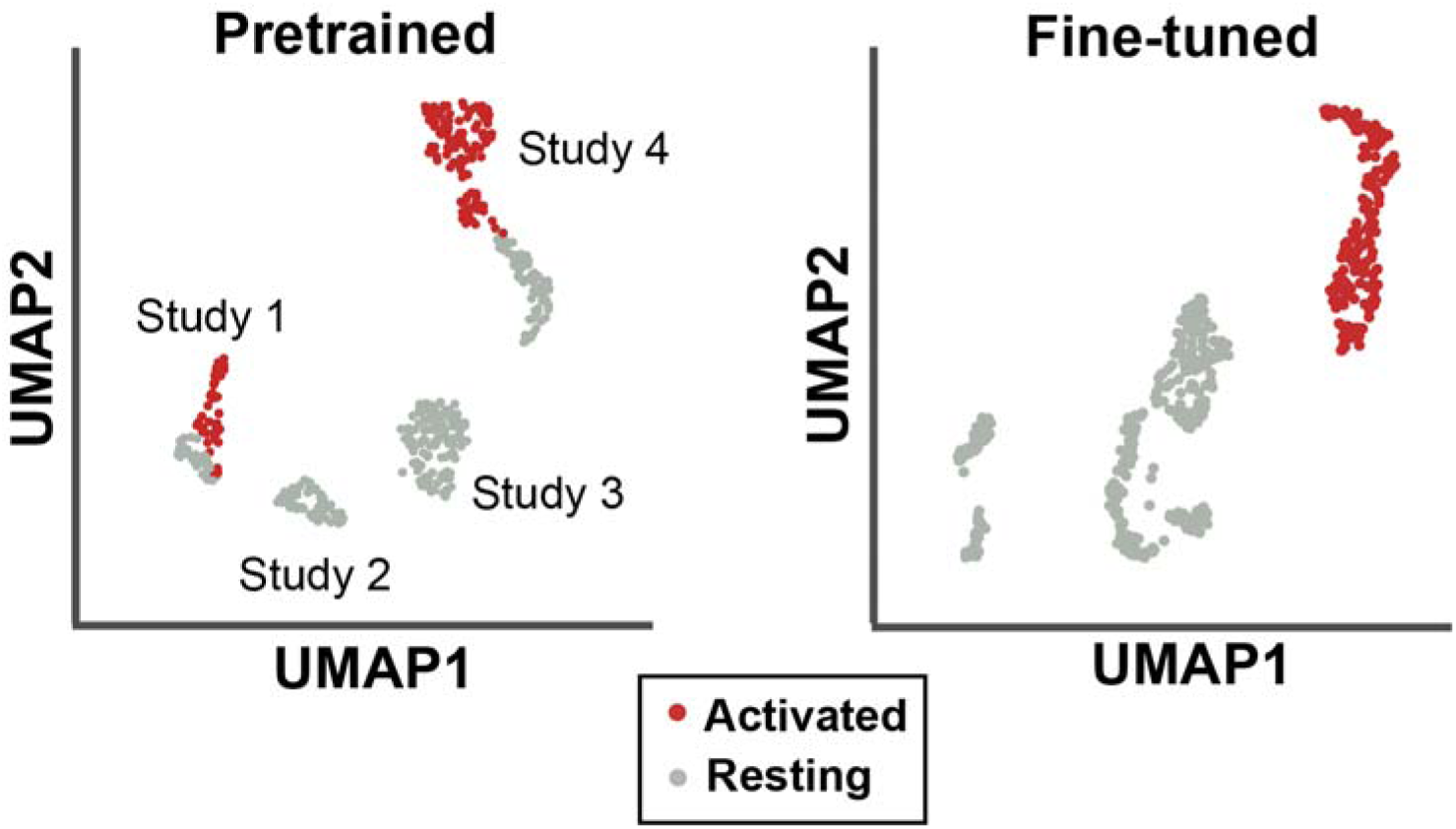
Embeddings before and after fine-tuning for T cell activation. UMAP of 512-dimension embeddings from Geneformer-30M-12L before and after fine-tuning using 4 studies of T cell activation for ∼20,000 T cells. Left shows pretrained embeddings, which cluster by study. Right shows fine-tuned embeddings, which cluster by activation status.

**Figure S2.**
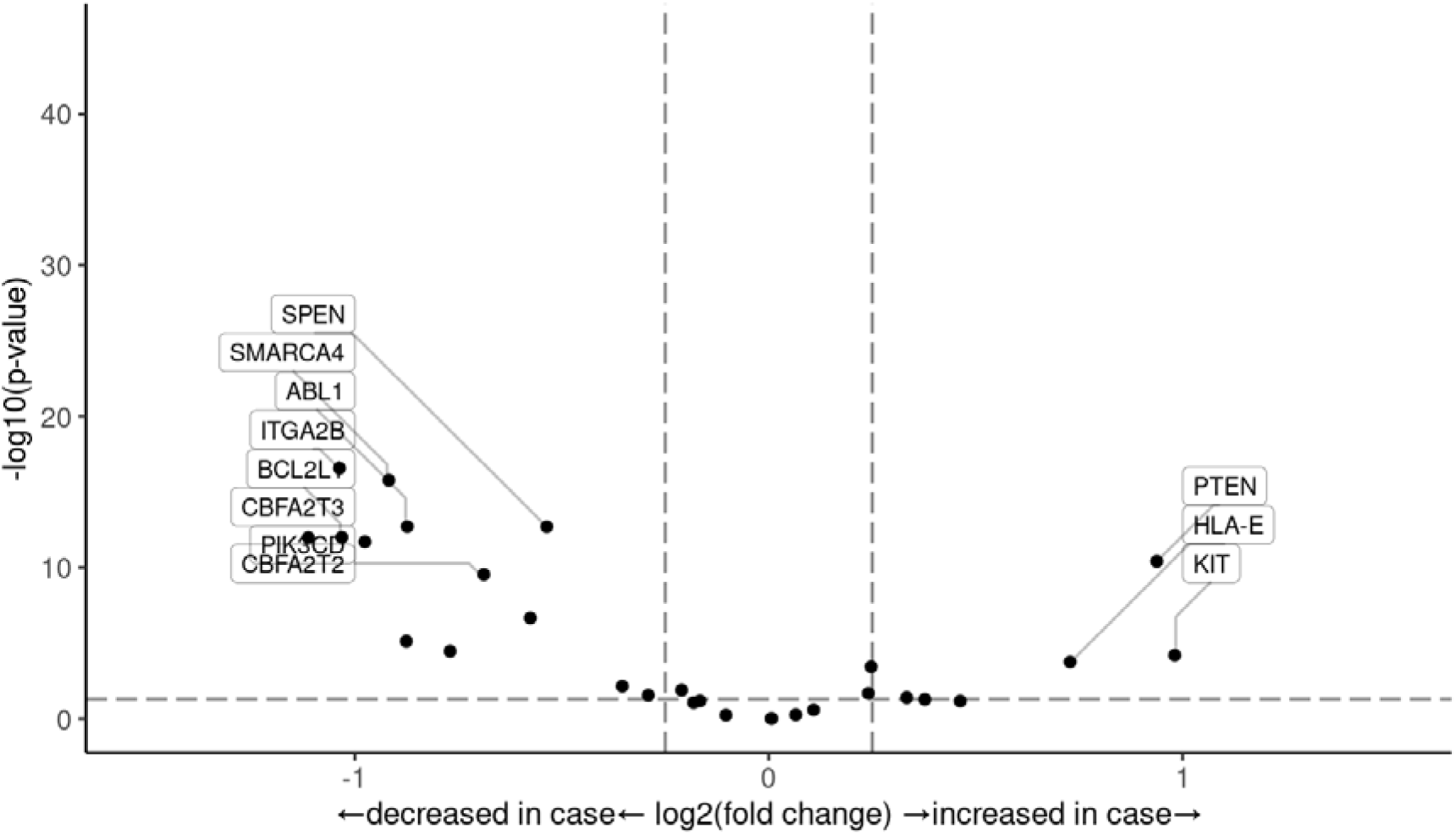
Differential expression of known RUNX1 targets between RUNX1-knockout hematopoietic stem cells (HSCs) and AAVS1-knockout HSCs. Most RUNX1 targets have a reduced expression in the setting of RUNX1-knockout.

**Figure S3.**
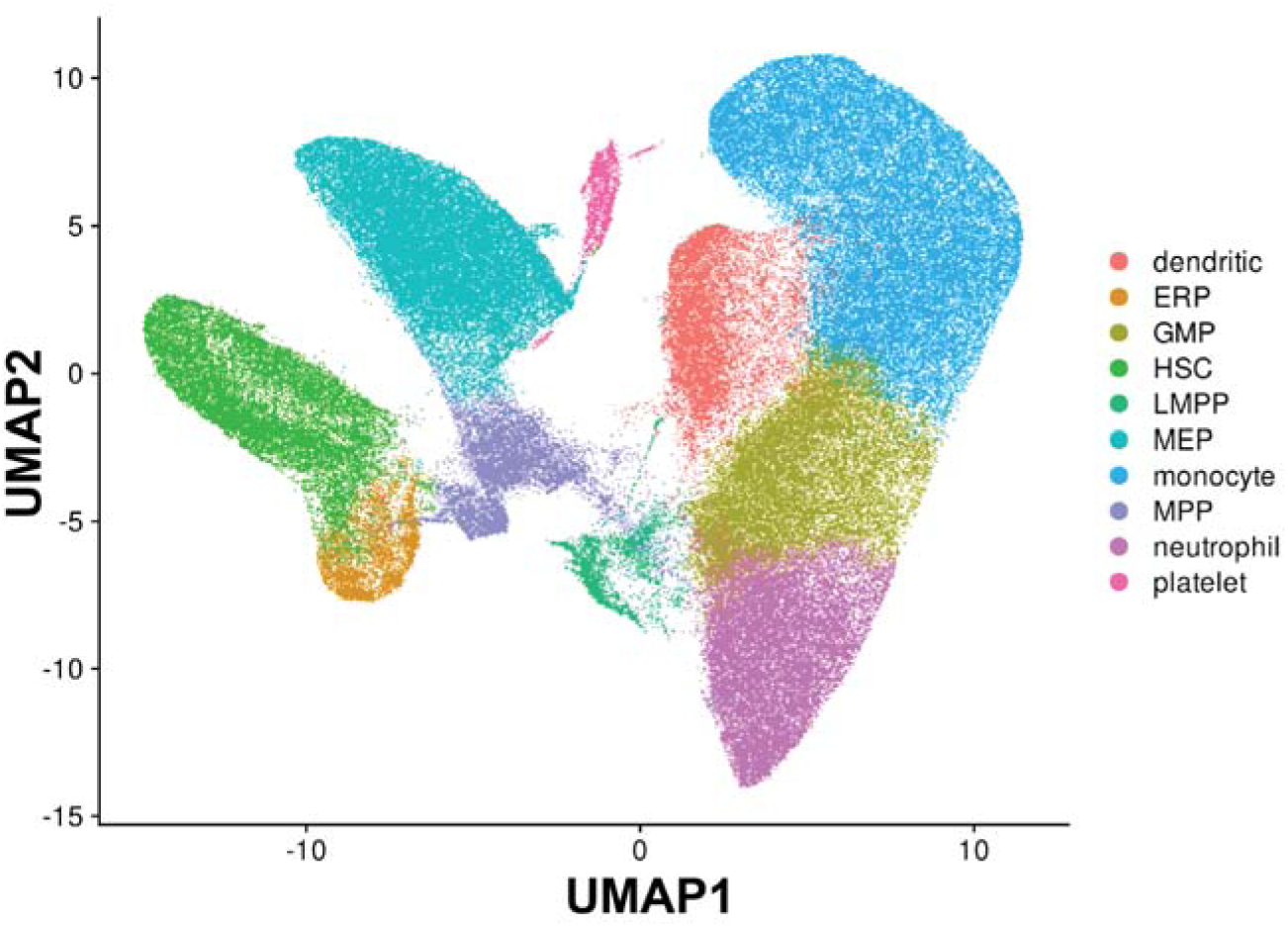
Single-cell RNA sequencing data generated for the *in vitro* model of *RUNX1*-knockout. Cell-type annotation identifies 10,243 hematopoietic stem cells (HSCs) which were then used for downstream perturbation analysis.

**Figure S4.**
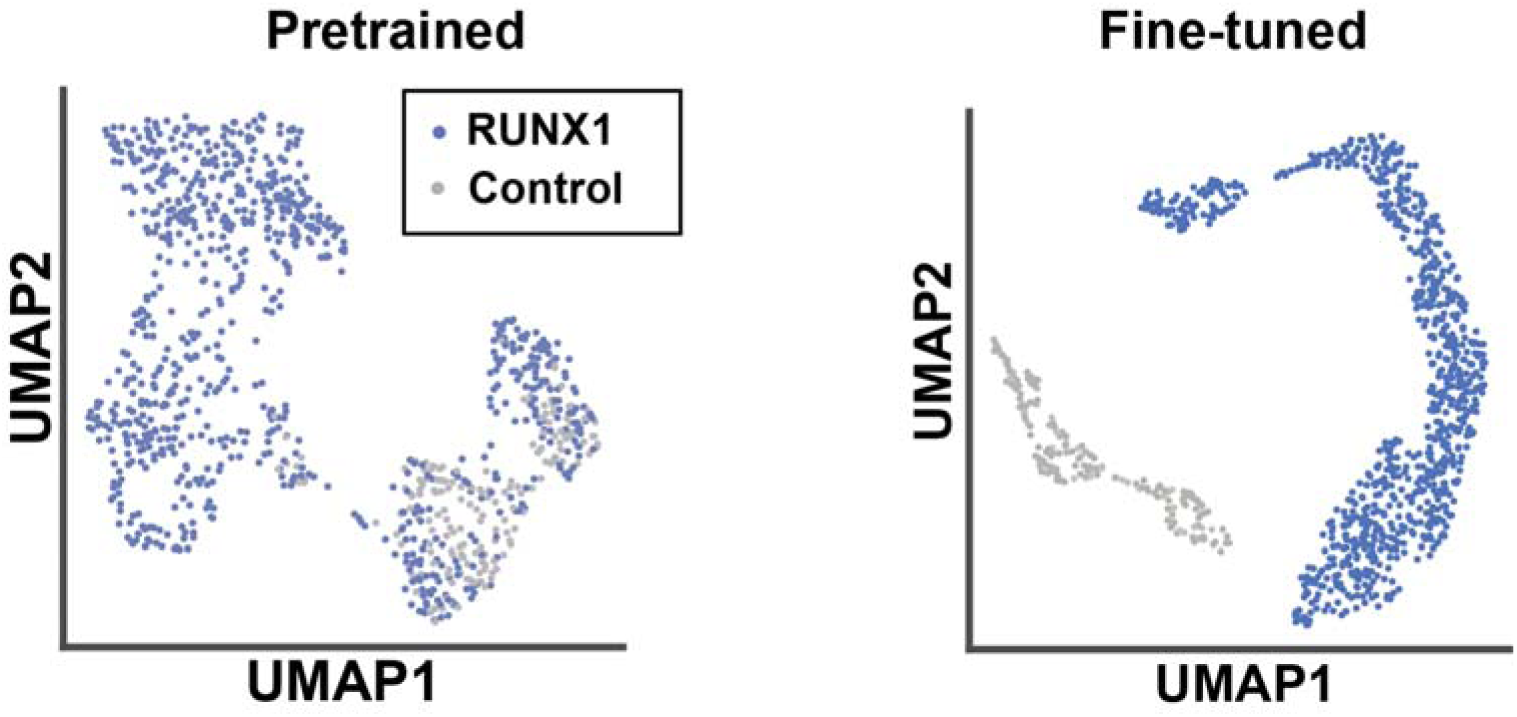
Embeddings before and after fine-tuning classified by sgRNA target (*RUNX1* or *AAVS1* control). UMAP of 512-dimension embeddings from Geneformer-30M-12L before and after fine-tuning 10,243 CD34+ hematopoietic stem and progenitor cells (HSPCs). Left shows pretrained embeddings. Right shows fine-tuned embeddings, which cluster by sgRNA.

**Figure S5.**
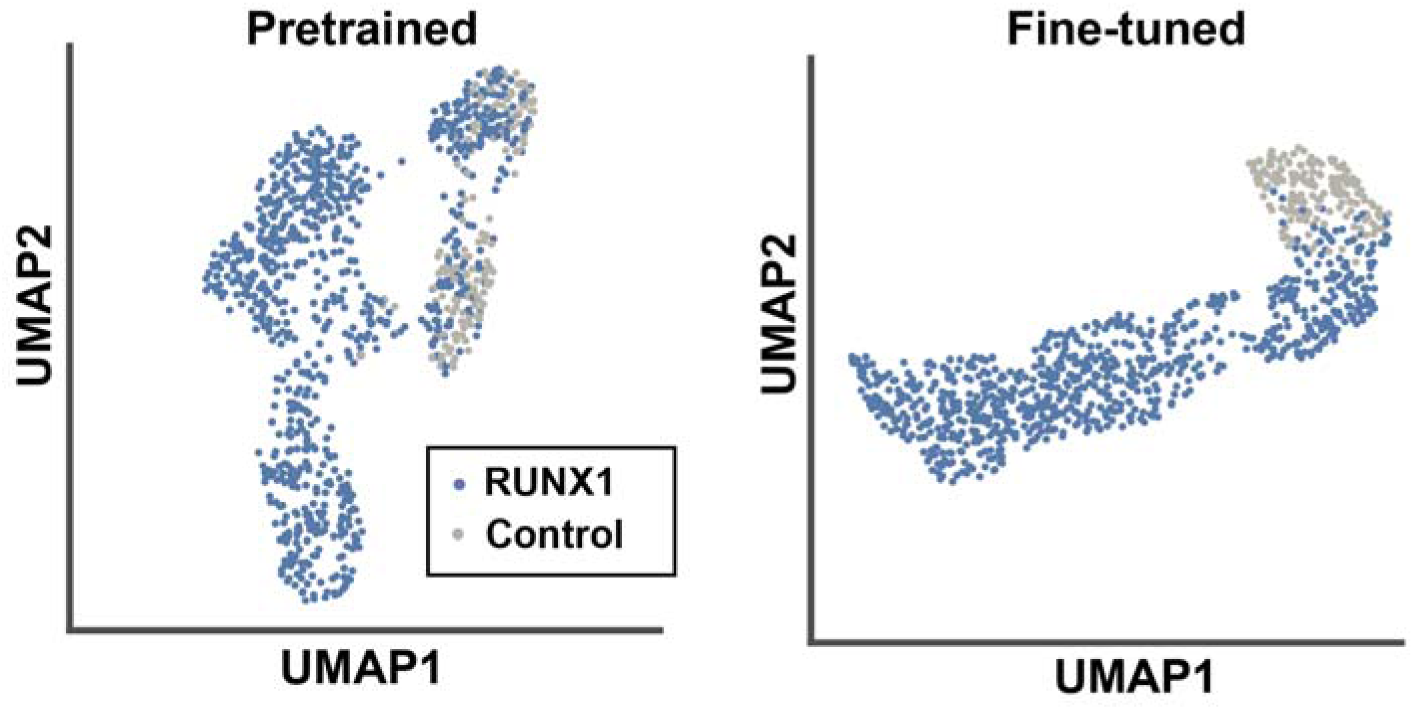
Embeddings before and after fine-tuning classified by sgRNA target (*RUNX1* or *AAVS1* control) including *RUNX1*-knockout cells having been exposed to small molecule perturbations. UMAP of 512-dimension embeddings from Geneformer-30M-12L before and after fine-tuning 9,806 CD34+ hematopoietic stem and progenitor cells (HSPCs).

## Methods

### Primary T cell activation training data

We used publicly available single-cell RNA sequencing data from 4 studies where T cells were either unstimulated or stimulated via CD3-CD28 beads or phorbol myristate acetate/ionomycin (PMA/ionomycin):

1. Kartha et al^15^ performed scRNAseq on resting and stimulated primary human peripheral blood mononuclear cells (PBMCs) from 4 donors. We included the scRNAseq data from 3,708 T cells which were unstimulated and 3,797 T cells which were stimulated with PMA/ionomycin along with Brefeldin A, a protein secretion inhibitor which allows us to isolate the primary effect of T cell activation. They were stimulated with PMA and Ionomycin calcium salt with or without Brefeldin A.
2. Cano-Gomez et al^16^ performed scRNAseq on primary T cells with and without anti-CD3/anti-CD28 beads in the absence of cytokines (Th0). CD4^+^ T cells were isolated from the PBMCs of 6 donors. T cells were then stimulated with anti-CD3/anti-CD28 human T-activator dynabeads. They profiled gene expression (RNA-seq) at 16 hours. A total of 5,269 resting T cells were profiled and 7,309 stimulated T cells were profiled.
3. Lawlor et al^17^ performed scRNAseq on PBMCs from 10 donors. PBMCs were cultured in control media (resting) versus plate-bound anti-CD3 + anti-CD28 (stimulated). 4,589 T cells were stimulated and 3,694 T cells were resting.
4. Szabo et al^18^ performed scRNAseq T cells, after CD3+ selection, from the blood of two deceased adult organ donors and two healthy adult volunteers with and without T-cell receptor stimulation using anti-CD3/CD28 T-activator for 16 hours. There were 9,147 resting T cells and 8,478 activated T cells.

For analysis with Geneformer, the single-cell RNAseq data was combined into a Seurat object and converted to a Loom object for tokenization. For DE analysis, Harmony was used to integrate the data from the four studies. Cells were grouped based on similarity of cell states using the Python package Metacell-2 with a targeted metacell size of 160,000 transcripts. Differential expression testing was performed using the Wald test for negative binomial regression through DESeq2 for genes that had at least 10 transcripts in at least 85% of metacells.

### Foundation model architecture and pretraining

We utilized Geneformer-30M-12L, a context-aware, attention-based deep learning model pretrained on approximately 30 million single-cell transcriptomes (Genecorpus-30M) as described by Theodoris et al.^5^ The model architecture consists of twelve transformer encoder units with an input size of 2048 and an embedding dimension of 256, four attention heads per layer, and a feed-forward size of 512. Each single-cell transcriptome was represented as a rank value encoding, where genes were ranked by their expression in each cell normalized by their median expression across the entire Genecorpus-30M. Pretraining was accomplished using a self-supervised masked learning objective, where 15% of genes within each transcriptome were masked, and the model was trained to predict which gene should be within each masked position using the context of the remaining unmasked genes. This pretraining strategy enabled the model to learn fundamental gene regulatory network dynamics in a completely self-supervised manner without requiring labeled data.

We utilized the *TranscriptomeTokenizer* class to convert our loom-formatted datasets into tokenized datasets compatible with Geneformer-30M-12L. We preserved relevant cell metadata (e.g., activation status, study source, cell type annotations, and count information). The tokenization process maintains the rank value encoding representation of each cell’s transcriptome, which prioritizes genes that distinguish cell states while deprioritizing ubiquitously expressed housekeeping genes.

### In silico perturbation prediction for T-cell activation

For predicting T-cell activation status, we fine-tuned the pretrained Geneformer-30M-12L model using publicly available scRNA-seq data from four independent studies where T cells were stimulated via CD3-CD28 beads or PMA/ionomycin, comprising a total of 37,991 cells (20,707 activated, 17,284 resting). The fine-tuning process was implemented using a custom classifier framework with a learning rate of 5×10^-5^, batch size of 6, gradient accumulation steps of 2, and linear learning rate scheduling with warm-up over 10 epochs. All 12 transformer layers were kept unfrozen during training to allow complete model adaptation to the T-cell activation task

After fine-tuning, we performed in silico perturbation (ISP) using the *InSilicoPerturber* class to systematically analyze the effect of genetic perturbations on T-cell activation state. For each gene in the GeneCorpus, we simulated both overexpression (by moving the gene to the front of the rank value encoding) and knockout (by removing the gene from the encoding). Cell embeddings were extracted from the final layer (layer 12) of the model, as these contain the most task-specific representations relevant to T-cell activation. We quantified the effects of these perturbations by measuring the cosine similarity between the original and perturbed cell embeddings, allowing precise determination of how each perturbation shifted cells toward either activated or resting states. The *InSilicoPerturberStats* class was used to process and analyze the large-scale perturbation data, calculating effect sizes and statistical significance for each gene perturbation.

### Evaluation of ISP for T cell activation with flow cytometry

We validated our predictions against orthogonal flow cytometry data from CRISPR activation and interference screens that measured IL-2 and IFN-γ production after CD3-CD28 stimulation in over 18,000 genes. This allowed systematic evaluation of ISP predictions compared to differential expression (DE) analysis using ground truth experimental data as a benchmark. To determine the minimum number of perturbation examples required for significant improvement, we systematically evaluated model performance with incremental additions of random perturbation subsets. Performance metrics including positive predictive value, negative predictive value, sensitivity, and specificity were calculated using flow cytometry data as ground truth to quantify the improvements gained through the closed-loop approach.

### Fine-tuning with T cell Perturbseq data

To enhance prediction accuracy, we developed a "closed-loop" ISP framework incorporating experimental perturbation data during model fine-tuning. We integrated both unperturbed T-cell transcriptomes and Perturbseq data from CRISPR activation and interference screens in primary human T cells. Critically, the Perturbseq data was labeled only with activation status rather than specific gene perturbation information, enabling the model to learn cellular responses to genetic perturbations while remaining agnostic to the specific targets.

This fine-tuning approach used the *BertForSequenceClassification* architecture with *DataCollatorForCellClassification* to handle batching and preprocessing of transcriptomic data. The model was fine-tuned to distinguish between activated and resting T-cell states using gradient-based optimization with early stopping based on validation performance. We systematically evaluated the minimum number of perturbation examples required for significant improvement by incrementally adding random subsets of perturbation data to the fine-tuning dataset, measuring performance metrics at each increment.

Following fine-tuning with the closed-loop approach, we performed ISP on all genes except those 75 genes included in the Perturbseq training data to avoid data leakage. The *EmbExtractor* class was used to extract and analyze cell-state embeddings, enabling precise quantification of cellular state shifts in response to in silico perturbations.

### In vitro model of RUNX1-Familial Platelet Disorder

Mobilized peripheral blood (mPB) CD34+ cells were purchased from Fred Hutchinson Cancer Center and introduced short indels using CRISPR-Cas9 with the ThermoFisher Neon NxT Electroporation system (1650V, 10ms, 3 pulses). Two days post-thaw, mPB CD34+ cells received ribonucleoprotein complexes with Cas9 (IDT Alt-R HiFi sp Cas9 Nuclease V3) plus sgRNA (IDT) at a 1:3.26 ratio towards *RUNX1* or *AAVS1*. Guide sequence was 5′-UACCUUGAAAGCGAUGGGCA-3′ for *RUNX1* as previously used by Fan et al^35^ and and 5’-GCCACTAGCCAGCCCGTCCG-3’ for *AAVS1* as previously described by Bak et al.^23^ Editing efficiencies were assessed using TIDE^36^ on PCR amplicons with primers targeting the *RUNX1*, which can be found in **Supplemental Table 6**. Cells were cultured for 9-15 days in CD34+ expansion media, containing the following reagents purchased from StemCell Technologies: SFEMII with 10% CD34+ expansion cocktail, 750nM StemReginin-1, 500nM UM729, and 20 U/mL of Penicillin/Streptomycin. Cells were kept at 150k-1M cells per mL.

### Single-cell RNA sequencing of HSPCs

For single-cell RNA sequencing analysis, mPB cells were cultured and harvested at various time points (7-13 days post-CRISPR editing). Cells were fixed using the Scale Biosciences fixation protocol found in standard methods or using the ScalePlex kit according to the manufacturer’s instructions. Fixed cells were counted and loaded at 20,000 cells per well into the Scale Biosciences platform. Single-cell RNA-seq libraries were prepared following the manufacturer’s protocol, and the quality of the libraries was assessed using Qubit fluorometry, qPCR, and Agilent BioAnalyzer. Libraries were sequenced on an Illumina NovaSeq X platform targeting 20,000 reads per cell.

### Differential expression of RUNX1-knockout and control HSPCs

Processing and analysis of scRNAseq data was performed primarily in R with the Seurat package. Doublets were identified with the Python package scrublet and removed. Cells with reads from fewer than 200 unique genes or fewer than 800 total transcripts were removed. A threshold was also set to exclude cells with greater than 25% mitochondrial transcripts, but capture of mitochondrial reads was negligible. Normalization and scaling was performed with SCTransform, followed by standard Seurat methods for dimensionality reduction and clustering. Top marker genes for each cluster were identified with the FindAllMarkers() function. Annotation of cell types was performed manually, based on marker genes. HSPCs were identified based on expression of *KIT*, *CD34*, and *CD38*.

Single-cell measurements were collapsed into metacells to address sparsity challenges. Cells were grouped based on similarity of cell states using the Python package Metacell-2 with a targeted metacell size of 160,000 transcripts. Cells that were delivered the sgRNA for *RUNX1* but clustered with the *AAVS1* cells were predicted to be unedited cells and were thus excluded from differential expression analysis. Differential expression testing was performed using the Wald test for negative binomial regression through DESeq2 for genes that had at least 10 transcripts in at least 85% of metacells.

### In silico perturbation prediction for RUNX1-knockout versus control

We fine-tuned the pretrained Geneformer-30M-12L model using scRNAseq from 10,000 HSPCs after CRISPR to predict *RUNX1*-knockout vs AAVS1 control status using the same process as for T-cell activation. After fine-tuning, we performed in silico perturbation (ISP) using the *InSilicoPerturber* class to systematically analyze the effect of genetic perturbations on cell state as described above.

### Drug screening and in silico perturbation using RUNX1-knockout HSPCs

We performed a high-throughput drug screen using HSPCs after CRISPR-Cas9 using an sgRNA targeting *RUNX1* as described above. Small-molecule inhibitors targeting predicted gene products were obtained from MedChem Express and the Vanderbilt High-Throughput Screening (HTS) core. Cultured mPB cells were treated with compounds at various concentrations for 24 hours, then fixed using Scale Biosciences fixation reagents for single-cell RNA sequencing. The small molecules are listed in **Supplemental Table 5**. HSPCs were identified during post-processing based on established marker genes, as described above. Then, we integrated both unperturbed HSPCs and HSPCs after chemical perturbation as data for fine-tuning the model, as was done with the Perturbseq T cell activation data above. Following fine-tuning with the closed-loop approach, we performed ISP as above, excluding the targets of the chemical perturbations that we used for fine-tuning. The *EmbExtractor* class was used to extract and analyze cell-state embeddings, enabling precise quantification of cellular state shifts in response to in silico perturbations.

## Data Availability

The single-cell RNA sequencing data for the in vitro model of *RUNX1*-FPD and the 4 T cell activation studies for T cell activation are available on Chan Zuckerberg Initiative’s CELLxGENE platform. The PerturbSeq data from Schmidt et al 2022 is available here: https://zenodo.org/records/5784651.

## Code Availability

Code for Geneformer-30M-12L is available at https://huggingface.co/ctheodoris/Geneformer. Code from this manuscript to implement in silico perturbation and closed-loop in silico perturbation available at: https://github.com/bicklab/geneformer. A tutorial to perform in silico perturbation with Geneformer is available at https://github.com/bicklab/geneformer.

## Acknowledgements

We would like to thank Argonne National Laboratory for computing resources, Angela Jones and the Vanderbilt VANTAGE sequencing core for single-cell RNA sequencing and Joshua Bauer and the Vanderbilt High-Throughput Screening Facility in the Vanderbilt Institute of Chemical Biology for assistance with our high-throughput drug screen.

We would like to thank the Chan Zuckerberg Initiative for funding this work as part of the Patient-Partnered Collaborations for Single-Cell Analysis of Rare Inflammatory Pediatric Disease. This project has been made possible in part by grant SCB-PPC-0000000092 from the Chan Zuckerberg Initiative DAF, an advised fund of the Silicon Valley Community Foundation. Y.P. and H.K.G. were supported by NIH grant T32 GM007347. J.V.A and A.C.P. were supported by NIH grant T32 GM080178. A.G.B. is supported by NIH grants DP5 OD029586, R01 AG088657, R01 AG083736, a Burroughs Wellcome Fund Career Award for Medical Scientists, the E.P. Evans Foundation, a Pew-Stewart Scholar for Cancer Research award, supported by the Pew Charitable Trusts and the Alexander and Margaret Stewart Trust, a Hevolution/AFAR New Investigator Award in Aging Biology and Geroscience Research.

